# ECS-based investigation of chloroplast ATP synthase regulation

**DOI:** 10.1101/2020.04.28.066100

**Authors:** Felix Buchert, Benjamin Bailleul, Pierre Joliot

## Abstract

The chloroplast ATP synthase (CF_1_F_o_) contains a specific feature to the green lineage: a γ-subunit redox domain which contains a cysteine couple and interacts with the torque-generating βDELSEED-loop. Based on the recently solved structure of this domain, it was proposed to function as a chock. *In vitro,* γ-disulfide formation slows down the activity of the CF_1_F_o_ at low transmembrane electrochemical proton gradient 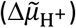. Here, we utilize *in vivo* absorption spectroscopy measurements for functional CF_1_F_o_ activity characterization in Arabidopsis leaves. The spectroscopic method allows us to measure the 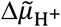 present in dark-adapted leaves, and to identify its mitochondrial sources. Furthermore, we follow the fate of the extra 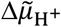 generated by an illumination, including its osmotic and electric components, and from there we estimate the lifetime of the light-generated ATP. In contrast with a previous report [Joliot and Joliot, Biochim. Biophys. Acta, 1777 (2008) 676-683], the CF_1_F_o_ γ-subunit exists mostly in an oxidized form in the dark-adapted state. To study the redox regulation of the CF_1_F_o_, we used thiol agent infiltration in WT and a mutant that does not form the γ-disulfide. The obtained 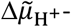 -dependent CF_1_F_o_ activity profile in the two γ-redox states *in vivo* reconciles with previous biochemical *in vitro* findings [Junesch and Gräber, Biochim. Biophys. Acta, 893 (1987) 275-288]. The highest rates of ATP synthesis we measured in the two γ-redox state were similar at high 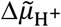. In the presence of the γ-dithiol, similar rates were obtained at a ~45 mV lower 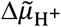 value compared to the oxidized state, which closely resembled the energetic gap of 0.7 ΔpH units reported *in vitro*.

## Introduction

During photosynthesis, the electron transfer is coupled to a movement of protons across the thylakoid membrane, generating an electrochemical proton gradient 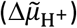 which consists of an osmotic component (the concentration gradient of hydrogen ions, ΔpH) and a membrane potential (ΔΨ). The chloroplast ATP synthase (CF_1_F_o_) catalyzes the conversion of the 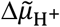 into the synthesis of ATP, subsequently used in the Calvin-Benson-Bassham cycle. The enzyme [recently reviewed in 1, 2] consists of two portions: a membrane-spanning F_o_ (subunits abb’c_14_) and a membrane-attached F_1_ (subunits α_3_β_3_γδε). In the chloroplast enzyme of vascular plants, during a 360° rotation of subunits γεc_14_ against the static subunits α_3_β_3_δabb’, 14 H^+^ are translocated along the electrochemical gradient while 3 molecules of ATP are synthesized. Flexibility in the peripheral stalk might redistribute torsional energy differences during the rotation cycle [3]. Perfect coupling is observed in isolated chloroplasts, i.e., H^+^ slip processes do practically not occur *in vitro* [4, 5, for a different view see Ref. 6]. Being reversible, CF_1_F_o_ catalyzes both the synthesis and hydrolysis of ATP and the reaction direction is determined by the extent of the 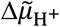 and the [ATP]/([ADP][P]) ratio.

The activity of the CF_1_F_o_ plays a central role in photosynthesis; it fuels the chemical phase of photosynthesis with ATP but can also participate in the regulation of the photochemical phase by modulating the osmotic component of the 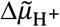. Indeed, the lumenal pH is involved in the regulation of the photosynthetic electron transfer at two levels. It regulates the photoprotective mechanism in photosystem II (PSII) and the turnover of the cytochrome *b_6_f* through the so-called photosynthetic control. Being at the crossroads of the dark and light phases of photosynthesis, the activity of CF_1_F_o_ needs to be fine-tuned and, although other regulations exist [7–9], two main regulatory mechanisms can modulate the rates of the CF_1_F_o_. The first one is relevant for F-ATP synthase in general which is a regulation by its substrate, the 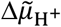. Above a light-induced threshold 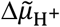, the transition of the inactive CF_1_F_o_ to a fully active form was observed in isolated chloroplasts [10, 11]. Several studies on chloroplasts and leaves demonstrated that each of the two components of the 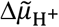 can activate the enzyme, provided that they allow the 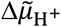 to reach this critical level [12–16]. We call this level 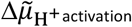 hereafter. Specific to CF_1_F_o_ in the green lineage is a second, “thiol modulation” or “redox regulation” mechanism which involves an insertion in the γ-subunit. Compared to cyanobacterial ancestors, the additional domain harbors a redox-active Cys couple which forms a disulfide in the dark. Formation of the γ-disulfide is, in part, mediated by a thioredoxin-like2/2-Cys peroxiredoxin redox cascade in Arabidopsis [17]. Dithiol formation *in vivo* is catalyzed by thioredoxin at moderate light intensities [18] and by NADPH thioredoxin reductase C in dim light [19, 20]. It was shown that the redox-active Cys is exposed to the solvent in the presence of a 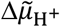 [21, 22], which could contribute to efficient dithiol formation in the presence of light [23, 24]. Moreover, various sites in CF_1_F_o_, some of them located in the γ-redox domain, were chemically labeled in the presence of 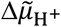 [21, 22, 25–28, reviewed in 29]. From a structural point of view, in the disulfide form, the γ-redox domain was found to act as an ATP hydrolysis chock that is vicinal to the torque-generating βDELSEED-loop [3, 30]. A particular γ-hairpin structure protrudes into the α_3_β_3_ hexamer and may interfere with efficient rotational catalysis. In the dark, when the 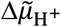 is low and the disulfide is present, the slowdown of ATP hydrolysis is believed to preserve high ATP levels [31]. However, point mutants of the redox-active Cys in *Arabidopsis thaliana* are viable [32]. Another mutant, termed *gamera*, remains in the γ-dithiol state in the dark by expressing a redox-insensitive γ-subunit isoform [33] and displays a stay green phenotype after several days in the dark [34]. It was suggested that altered pH-dependent protein import might be, in part, responsible for the phenotype. Several γ-/β-subunit interactions, possibly the γ-hairpin structure as well, seem to be modulated by the γ-redox state [35]. Taken together, this suggests that CF_1_F_o_ experiences intertwined 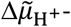 and redox-dependent structural changes. Junesch and Gräber demonstrated *in vitro* that the dependency between the redox and 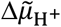 regulations of CF_1_F_o_ stems from a lower 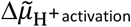 threshold in presence of the γ-dithiol [36], yielding half-maximal rates ~0.7 ΔpH units apart.

The knowledge accumulated through *in vitro* studies has not yet been confirmed *in vivo,* which is the main aim of this work. Because the CF_1_F_o_ activity is both 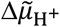 and redox regulated, the study of its redox regulation requires the *in vivo* measurement of both CF_1_F_o_ activity and 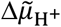 extent. The first can be measured via routinely used techniques. Since photosynthetic membranes are spectroscopic voltmeters, the ΔΨ can be probed linearly by ECS, the electrochromic shift of photosynthetic pigments [37, 38, reviewed in 39]. Following a light-induced membrane energization in dark-adapted samples, the ΔΨ (and ECS) decay is tightly linked to H^+^ translocation via CF_1_F_o_. ECS decay kinetics following a single-turnover laser flash are therefore commonly used to probe CF_1_F_o_ activity *in vitro* [e.g., Ref. 11] as well as *in vivo* [e.g., Refs. 33, 40]. In contrast, the measurement of the 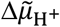 is a methodological bottleneck but a protocol was proposed previously [16] whose principle can be explained through a simple analogy. The water volume contained in an opaque bottle cannot be read, but its empty volume can be easily measured as the volume of water that needs to be added before it spills out (when the maximal countenance capacity is reached). Even if the maximal countenance capacity of the bottle remains unknown, the water volume can be given in relation to this reference/leak value. The same applies for the measurement of the electrochemical proton gradient in the dark. Joliot and Joliot have shown that a threshold of electrochemical proton gradient exists, corresponding to the leak of protons through the thylakoid membrane *in vivo* 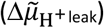. It is identified as the maximal ECS that can be sustained by a short illumination with very strong light. The 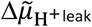 is a constant in a given photosynthetic material and can therefore be used as a “maximal countenance capacity” reference [16]. In brief, the ECS-based protocol to measure the extent of 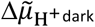 consists in energizing the membrane with a very strong light pulse and probing the ECS increase until the 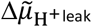 is reached (i.e., measuring the “empty volume”). This method was used previously with success [16], but some of the implicit hypothesis of this method deserved to be further tested: (i) The increase of ECS (which probes ΔΨ) reflects the increase of 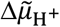 or, in other words, there is no significant ΔpH generation during the short saturating pulse. Moreover, since the saturating pulse surpasses 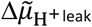, the leak process *in vivo* can either occur through 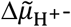-regulated H^+^ channels or via CF_1_F_o_. (ii) The method, tested only on dark-adapted leaves, remains valid whatever the redox state of the γ-subunit domain. In this work, we demonstrate that those hypotheses are respected making this method valid to assess the 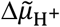 in both the reduced and oxidized states of CF_1_F_o_. This allowed us to confirm that CF_1_F_o_ is mostly in the oxidized state in dark-adapted leaves (in contrast with a previous report [16]) and to validate *in vivo* the model of Junesch and Gräber. Our results show that the 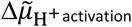 of the disulfide-containing CF_1_F_o_ was ~45 mV lower and, based on the CF_1_F_o_ activity profile in leaves infiltrated with a thiol agent, the efficiency of the chemical reduction of the γ-subunit was a little less than 50%. Finally, as a further improvement of the method, we *first* utilized the fact that the 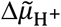 is in equilibrium with the [ATP]/([ADP][P]) ratio to follow the lifetime of ATP after a light perturbation, and *second* determined the 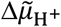 partitioning into the ΔpH and ΔΨ components when incrementing illumination time.

## Material and Methods

### Absorption changes

The experimental setup was as described previously [16]. Absorption changes were measured using a JTS spectrophotometer (Biologic). Pulses of saturating light are provided by LEDs peaking at 630 nm (~ 2500 μmol photons m^-2^ s^-1^), unless otherwise stated. This light irradiance corresponds to ~2000 photons absorbed per second per PSI or per PSII, based on the measurement of changes in the membrane potential (ΔΨ) according to the method described previously [16].

### Correction and normalization of ECS signals

The light-induced membrane potential changes (ΔΔΨ) were measured through the absorption changes at 520 nm (reflecting mostly ECS) for experiments with short pulses (less than 20 ms), and 520 nm-546 nm for longer pulses (Fig 2). Indeed, subtracting 546 nm allows to correct for the contribution of cytochromes, P700 and scattering. However, the comparison between corrected data (520-546) and uncorrected data (520) shows that the correction was not necessary for short pulses (Fig S1). To express the measured ECS signals in charge separations per photosystem I (PSI), we calibrated the ECS signal by the ECS increase corresponding to one charge separation per PSI. For that, we divided all measured ECS signals in this work by ½ the ECS increase following a single-turnover flash, provided by a dye laser at 690 nm pumped with a Nd:Yag laser. Since all photosystems will generate one charge separation upon such a flash, the ECS measured 250 μs after the flash is proportional to the number of active PSI + PSII. Assuming equal concentrations of both photosystem reaction centers, half this signal is equal to the sole contribution of PSI.

### Reproducibility

The kinetics of ECS rise and decay were reproducible between different leaves of different batches, but it was not the case of the extent of the ECS. This is because its amplitude depends on the density of reaction centers and ECS probes per leaf area, which slightly varied in different cultures. For this reason, for each experiment where the amplitude of ECS was probed, we compared samples from the same batch of culture.

### Chemicals and Inhibitors

We used chemical compounds by vacuum infiltration since the method did not interfere with the ΔΨ increase and decay kinetics as compared to non-infiltrated samples (see Fig S2). All chemicals were purchased from Sigma-Aldrich. Tris(2-carboxyethyl)phosphine hydrochloride (TCEP) was dissolved in water at 0.3 M and aliquots, adjusted to pH 7.0, were frozen until further use. Solutions of antimycin-A (AA, 40 mM), nonactin (40 mM), nigericin (Nig, 10 mM) and Carbonyl cyanide 4-(trifluoromethoxy)phenylhydrazone (1 mM) were dissolved in ethanol and usually served as 1000x stocks that were stored at −20°C.

## Results

### 1. Membrane potential decay measurements after saturating light pulses

In order to quantify the CF_1_F_o_ performance in γ-disulfide- and γ-dithiol-promoting conditions, we will first elaborate on an improved protocol which is based on a previous study [16] and allows to measure the 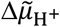 already present in dark-adapted leaves 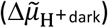. The method is based on the kinetics of the ECS and measures the ΔΨ changes (ΔΔΨ) induced by a short saturating light pulse, when compared to the baseline ΔΨ prior to the pulse (Fig 1). We make the reasonable hypothesis that short pulses (< 20 ms) induce exclusively ΔΔΨ and that changes in the ΔpH can be neglected [16] due to the high buffering capacity inside the lumen [41]. According to this hypothesis (tested in section 2), ECS changes which are strictly speaking proportional to ΔΔΨ, equally reflect changes in the electrochemical proton gradient 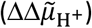. As explained before [16], since the CF_1_F_o_ is reversible and outpaces slow passive ion movements across the membrane, the 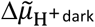 and the [ATP]/([ADP][P]) ratio reach a thermodynamic equilibrium in dark-adapted leaves, which make the 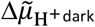 dependent on the physiological conditions. Indeed, thanks to an ancient nucleotide import machinery, the chloroplast ATP level is itself strictly dependent on the metabolic coupling with mitochondria [16, 42–46].

**Fig 1:**
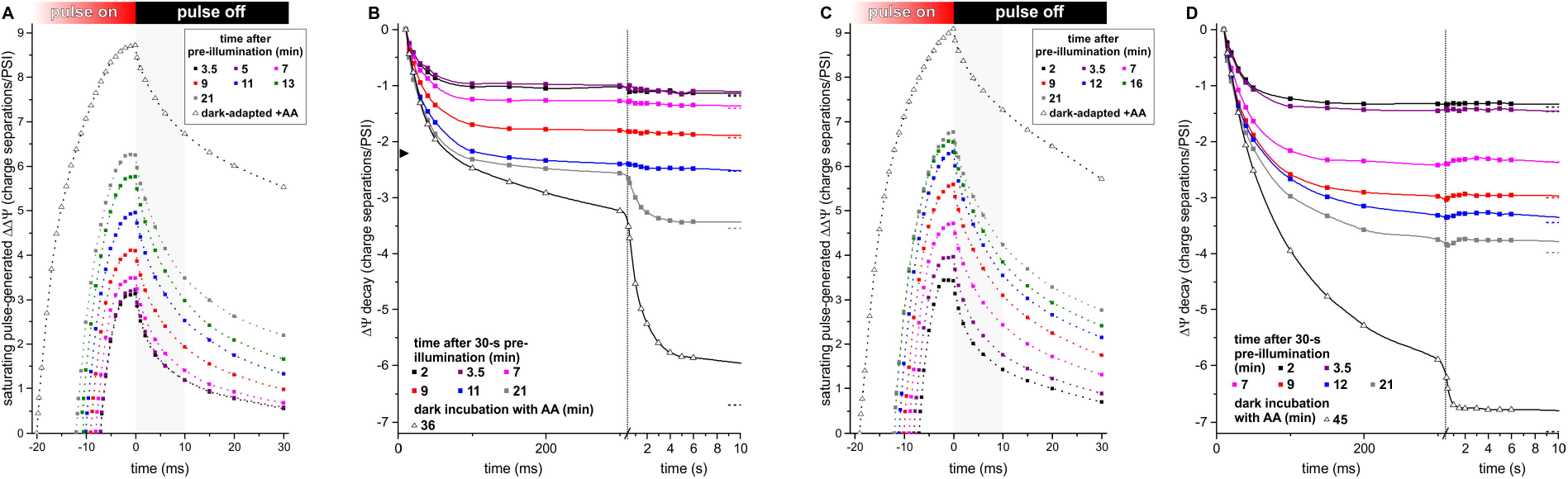
Pulse-induced ECS kinetics are shown in water-infiltrated leaves in the absence (A-B) or presence (C-D) of the mild reductant TCEP at 50 mM. Dark-adapted leaves were illuminated for 30 s (green-orange LED, 1000 μmol photons m^-2^ s^-1^), followed by a dark re-adaptation for several minutes (indicated in the legend). +AA: infiltrated with 40 μM antimycin-A without illumination. All the measurements were made on different leaves from the same plant. In panels A and C, the dark-adapted (baseline) ECS level was arbitrarily set to 0 and only the ECS kinetics during the pulse and in the initial relaxation phases are shown. In panels B and D, the ECS_10ms_ values of all curves were set to 0 and only the ECS relaxation in the dark is shown (the value reached after 1 min relaxation is indicated with three dots on the right). The transition between the fast and slow relaxation phases 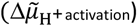 is indicated in panel B by an arrowhead.

Fig 1A illustrates the influence of the baseline 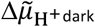 prior to the pulse on the amplitude and kinetics of the pulse-induced ECS (or 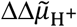). WT leaves were infiltrated with water and a low [ATP]/([ADP][P]) ratio was obtained under impairment of the mitochondrial respiration upon the addition of antimycin-A (AA), an inhibitor of the cytochrome bc1 (open symbols in Fig 1). On the contrary, high [ATP]/([ADP][P]) ratios were reached after a 30-s illumination (closed symbols). In all curves of Fig 1A, the ECS reached a plateau during the pulse 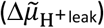 followed by a fast decay phase. Both the plateau and the amplitude of the fast phase depend on the light intensity (Fig S3). Such a fast phase has been observed in spinach [16] and is attributed to H^+^ leaks that are triggered above a critical 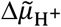 potential 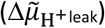, which is an absolute reference of the electrochemical proton gradient in a given material *in vivo.* However, after 10 ms decay, the ECS decay kinetics becomes independent of the light intensity (the leak process contributions is over at this time, Fig S3) and reflect the H^+^ release from the lumen via CF_1_F_o_ that is associated with ATP synthesis [16]. The ECS reached 10 ms after the end of the pulse (ECS_10ms_) is therefore independent of pulse intensity (see Fig S3), and also independent of the physiological conditions preceding the pulse [16]. Thus, the ECS_10ms_ will be used hereafter as the experimental reference of 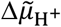 and ECS data in the following will appear in negative values (ECS - ECS_10ms_; Fig 1B) and correspond to the “empty volume” of the opaque bottle analogy. The light intensity of the pulse will be kept constant throughout the study (see Material and Methods).

The amplitude of the ECS increase was higher in the presence of AA than after the pre-illumination (Fig 1A). Given that the reached 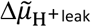 at the end of the pulse is an absolute reference of 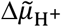, the higher pulse-induced ECS in the presence of AA did not reflect a higher 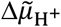 at the end of the pulse in those conditions. It rather reflected a lower baseline 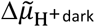 before the pulse, which was revealed after rescaling to our reference ECS_10ms_ (−6.7 vs. >−3.5 charge separations/PSI, Fig 1B). The ECS amplitudes during the pulse depended on the 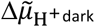 which is a pedestal onto which an extra 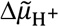 is superimposed until the total 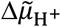 reaches 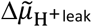 leak [16]. After the 30-s illumination, the increase of the amplitude of the pulse with the time in darkness (Fig 1A), again, indicated the continuous decrease of the 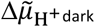 (Fig 1B) and reflected the consumption of the extra ATP produced during the 30-s illumination. According to Fig 1B and a previous report [16], the ECS decay kinetics after the pulse displayed three phases. A first phase, completed in 10 ms, was associated with H^+^ leaks (Fig 1A) as discussed before and will be discarded from now on. A second phase was completed in ~100 ms (phase 2) and was linked to ATP synthesis. Then, a multiphasic slow decay (phase 3), completed in ~1 minute, was observed only once the 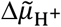 fell below a threshold (at the end of phase 2, ~100 ms). This threshold corresponds to the 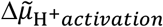 described before [11–16] and represented a second reference level [15], about 2 charge separations/PSI below ECS_10ms_ (arrowhead in Fig 1B). Phase 3 indicated a slowly active CF_1_F_o_, or, alternatively, a small CF_1_F_o_ fraction staying fully active.

We first decided to check whether this quantitative measurement of the 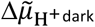 remained valid regardless of the redox regulation mechanism. Figs 1C and 1D show similar experiments on leaves infiltrated with a thiol reductant, TCEP. Furthermore, the samples were in the same physiological conditions as in Figs 1A/B (AA-treated or pre-illuminated samples). In all cases, the ECS generation and the decay in the first tens of ms after the end of the pulse resembled the untreated sample. As in water-infiltrated leaves, AA-treated leaves in the presence of TCEP showed a lower 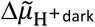 and illumination increased this pedestal (dashed lines in Fig 1D), reflecting ATP accumulation during preillumination. The 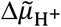 decay kinetics were always monophasic; we did not observe the slow phase of ECS decay (absence of phase 3 in Fig 1D). This is true even in conditions where 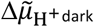 was below the 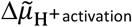 of Fig 1B, i.e. when dark re-adaptation after the 30-s illumination was longer than 9 min or in AA-treated samples. This observation was in agreement with the higher CF_1_F_o_ activity measured *in vitro* upon cleaving the γ-disulfide by chemical thiol reductants, at low 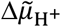 only [36]. A quantitative analysis will be given in section 4, demonstrating how the effect of the γ-redox state influences the 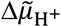 dependence of CF_1_F_o_ activity.

### 2. Relative extent of ΔΨ and ΔpH induced by short light pulses

For the interpretation of the previous results regarding the evolution of 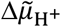, we followed the opaque bottle analogy and have made the hypothesis that the experimental reference of the short saturating light pulses, ECS_10ms_, served as our 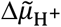 reference. This implied that the ΔpH that was generated until reaching ECS_10ms_ was insignificant, and only ΔΨ increased during the short saturating light pulses. To test our hypothesis, we first obtained a non-perturbed ECS_10ms_ reference from dark-adapted (for several minutes) leaves by applying a short 12-ms probing pulse (dark line, Fig 2A). Then, we applied saturating light pulses of variable duration to the dark-adapted samples and probed, after a relatively short dark period of 15 s, the extra ΔΨ (ΔΔΨ) that was sustained. The ΔΨ was monitored throughout the measurement and ΔΔΨ was calculated as the difference between the dark-adapted ECS signal before the first light perturbation and the ECS signal 15-s later (Fig 2A). The extra 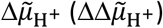 that was present 15-s after the first light perturbation was obtained, again, from a 12-ms probing pulse (i.e., from ECS_10ms_). Especially after prolonged light pulses, the ECS_10ms_ was below the non-perturbed ECS_10ms_ from dark-adapted material. We attributed this lowering of our electrogenic 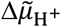 reference to the extra ΔpH (ΔΔpH) that was present after 15 s dark. A duration of 15 s dark was necessary in this experiment to allow a partial relaxation of the electron carriers toward the dark-adapted state. Indeed, at the end of the first light pulse, the photochemistry of both photosystems was low due to the absence of acceptors (oxidized plastoquinones or oxidized ferredoxins) or donors (reduced P700). If the second light pulse was applied too early, photochemistry was insufficient to reach the 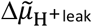 which made measurement of the 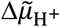 impossible.

**Fig 2:**
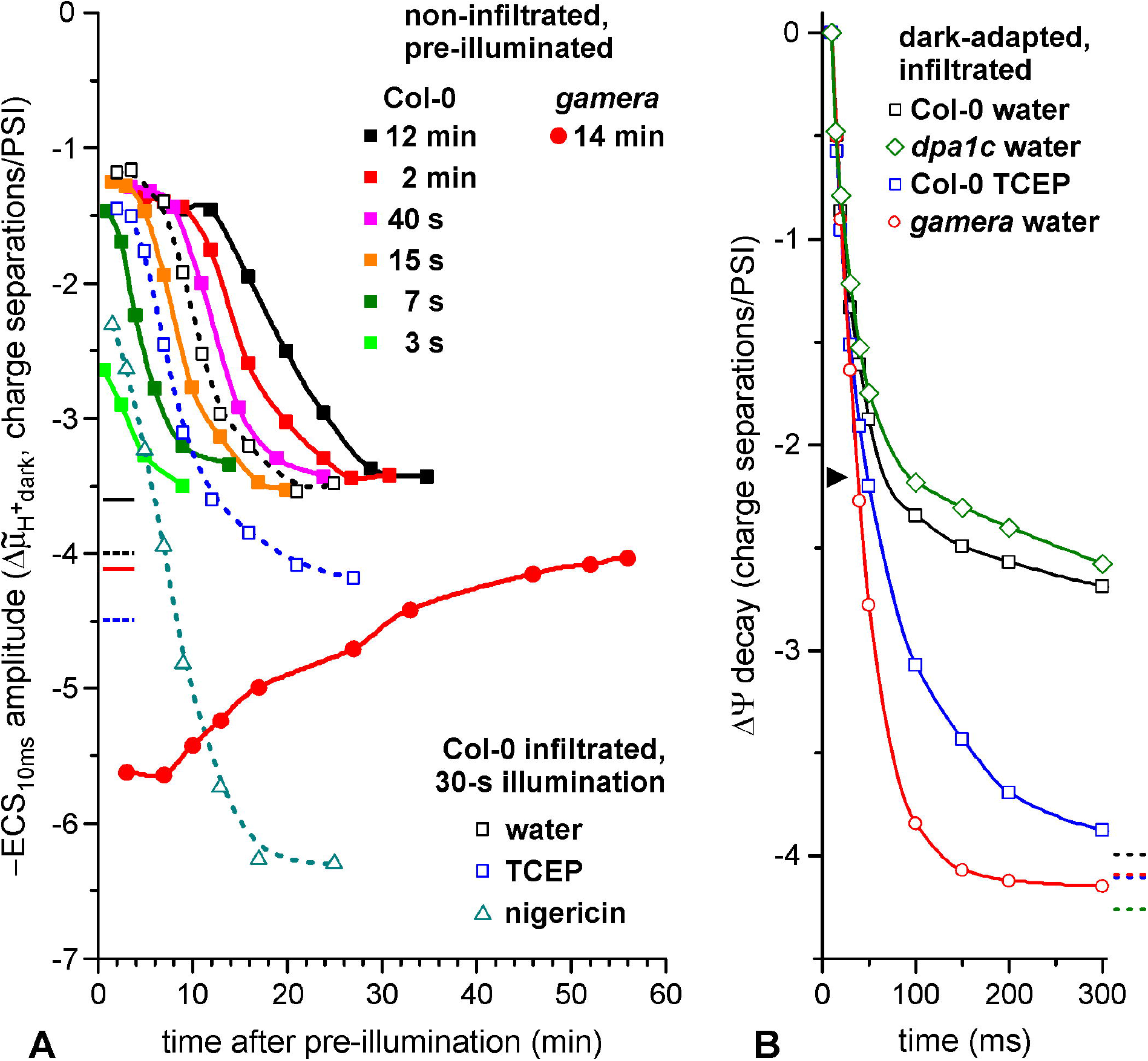
The estimation of ΔΔΨ, ΔΔpH and 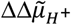 generated by an illumination is shown. The parameters were measured 15 s after a saturating pulse of various duration. (A) Exemplary ECS measurement in the presence of nigericin (open symbols light on, closed symbols darkness) measured as the difference between the absorption changes at 520 nm and the one at 546 nm (see Methods). The leaf was illuminated with a first pulse of 500-ms (*t* = −15 s), then ECS was followed for 15 s in the dark and a 12-ms probing pulse was used to determine the new ECS_10ms_ reference (the probing pulse ended at *t* = 0 s). For the sake of visualization, the dark-adapted ECS decay was shifted on the x-axis by 3 ms. (B) and (C) The initial phase of the ΔΨ decay after the 12-ms pulse is shown for leaves infiltrated with (B) water and (C) nigericin. The ΔΔΨ at the end of the 15-s dark period is indicated by horizontal dashed lines. The ΔΔpH corresponds to the difference between the ECS_10ms_ after the saturating pulse and the one of a dark-adapted leaf (dark closed squares). (D) and (E) The relationship of ΔΔΨ, ΔΔpH and 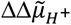 with the duration of the first light pulse are shown for water (D) and nigericin (E) infiltrated leaves. All the measurements in this figure were performed with leaves from the same plant.

We followed those parameters in water-infiltrated and in nigericin-treated leaves for different durations of the first pulse (10 ms to 1 s), given 15 s before the probing 12-ms pulse. For every first pulse duration, the ECS_10ms_ upon the second pulse was smaller than in the corresponding dark-adapted sample (Fig 2B). This indicated that a significant amount of extra ATP, generated during and after the first illumination, was still present after 15 s dark and increased of the 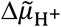 (more precisely, ΔΔpH) compared to the dark-adapted level. There was a significant change in the lowered ECS_10ms_ between the 100-ms and 500-ms light perturbation. In water-infiltrated samples, the ΔΔΨ at 15 s after the first perturbation (dashed line in Fig 2B) was significant, up to 0.4 charge separation/PSI. The ΔΔΨ appeared to stagnate upon light perturbations longer than 100-ms.

We used the ionophore nigericin (Nig), which exchanges protons with K^+^, and it is important to note that Nig also collapsed the proton motive force for mitochondrial ATP synthesis and thus interfered with metabolic coupling between the organelles. Accordingly, Nig infiltration resulted in a collapse of 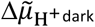 in the chloroplast (within ~15-30 min incubation, see Fig S4 and discussion). Here, Nig was used as a proof of principle to favor ΔΔΨ at the expense of ΔΔpH. Nig-infiltrated samples were analyzed in Fig 2C and the extent of the lowered ECS_10ms_ as a function of the first light pulse duration (ΔΔpH) showed similar developments as the controls in Fig 2B. However, the ΔΔΨ after 15 s was always higher than 1 charge separation/PSI, even for the shortest durations of the light pulse (dashed line in Fig 2C). The probed 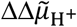 stored 15 s after a 10-ms pulse was 0.4 and 1.4 charge separations/PSI in control and Nig samples, respectively. For a 1-s pulse, those values reached 1.1 and 2.8 charge separations/PSI for control and Nig samples, respectively. In water-infiltrated samples (Fig 2D), the dependency of ΔΔpH and ΔΔΨ with respect to the first pulse duration indicated that short light pulses (100-ms or below) induced mostly ΔΔΨ whereas ΔΔpH dominated after longer light pulses (500-ms or 1-s). In contrast (Fig 2E), presence of Nig promoted a very strong ΔΔΨ which was present 15 s after the first light pulse, whereas the amplitude of the ΔΔpH was a lot smaller regardless of the pulse duration.

The fact that short pulses (below 100 ms) generated only small changes in ΔpH validated our initial hypothesis. The extent of ΔpH was smaller than 0.1 charge separation/PSI. This indicated that the ECS pulse method allowed the quantitative measurement of 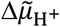 with a potential error of 0.1 charge separation/PSI only. The higher 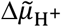 storage after 15 s dark in the Nig samples was expected, simply because the relaxation of the pulse-induced 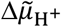 was slower when the 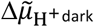 was below the 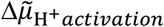, as we described before in the case of a AA treated sample (Fig 1B). Compared to AA, the uncoupling by Nig was more efficient in collapsing 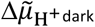 but, like other uncouplers, it modified the different phases of ECS decay kinetics (Fig S4, see also discussion). Therefore, the use of AA will be preferred instead of Nig in the following sections when collapsing the 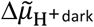 will be needed, because AA preserves membrane integrity.

### 3. What is the 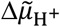 /ATP lifetime after an illumination?

In the previous sections, we have shown that our methodology allows quantitative measurements of the 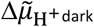 (relative to the 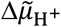 reference given by ECS_10ms_) provided that the saturating light pulse is (i) strong enough to reach the 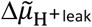 (which is valid for the 2000 photons/PSI/s light irradiance used here) and (ii) short enough to avoid the generation of ΔpH (which is valid for pulses shorter than 100 ms). In this section, we use this method to follow the fate of the 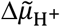 after an illumination by supposing that the thermodynamic equilibrium is met during a dark period before the ECS measurement and, as discussed before, following 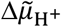 is equivalent to following the [ATP]/([ADP][P]) ratio. In Fig 1B, we measured the 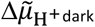 at different time in darkness following a 30-s illumination in water-infiltrated leaves. In Fig 3A, we reported the values of the 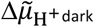 as a function of the time in darkness. In the water-infiltrated leaf, the 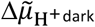 after the 30-s illumination was ~-1.2 charge separations/PSI relatively to the 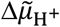 reference (ECS_10ms_), a value significantly higher than the dark-adapted one (~-4 charge separations/PSI, black dashed line). This value remained constant for 5 minutes following the illumination and, after this lag, relaxed to the dark-adapted value in ~30 minutes (Fig 3A). As discussed before, this reflected the build-up of ATP during illumination, and the subsequent slow consumption of this extra ATP. This consumption may be associated with slow passive H^+^ leaks through the membrane and/or with ATP-consuming reactions occurring in the chloroplast stroma. When leaves were infiltrated with Nig and illuminated for 30 s (Fig 3A), the 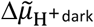 reached a lower level (−2.3 charge separations/PSI after 2 min dark) and relaxed to an even lower value within ~15 minutes (−6.3 charge separations/PSI). This low value in a dark-adapted sample reflected the fact that the 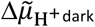 is suppressed due to the absence of mitochondrial ATP production in darkness, as discussed before. The ATP generated in the chloroplast during the illumination might be consumed more rapidly in the presence of Nig, probably because the metabolic needs cannot be met without mitochondrial ATP production.

**Fig 3:**
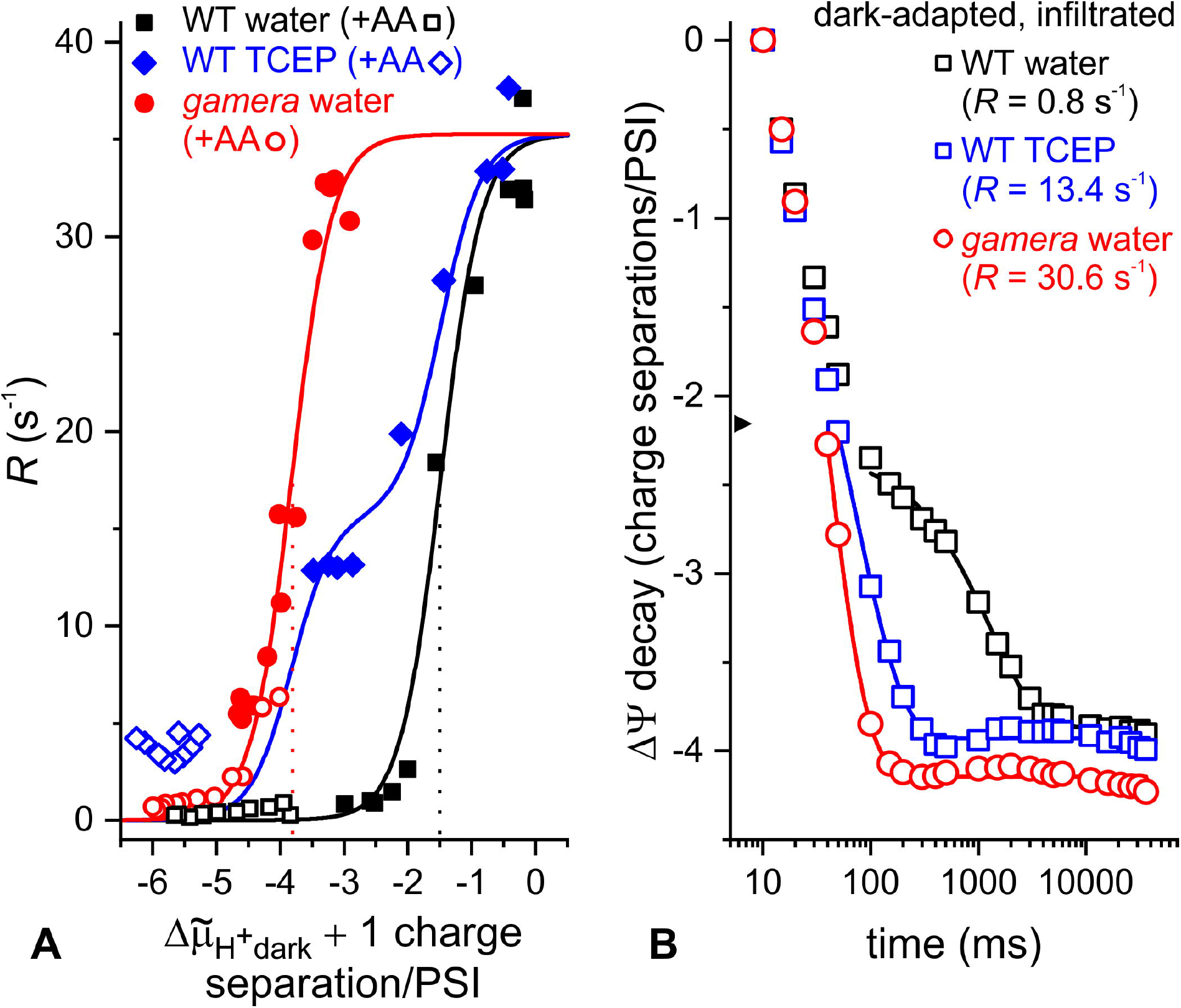
Light-induced ATP lifetime measurements and membrane potential (ΔΨ) decay kinetics in various Arabidopsis genotypes are shown. (A) The effect of the γ-disulfide bond reduction on ATP accumulation and relaxation after an illumination of different durations is displayed. The 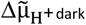 values are relative to the ECS_10ms_ and horizontal lines refer to fully dark-adapted, non-infiltrated (solid) and infiltrated (dashed) samples. The water-/TCEP-infiltrated WT samples were from the same plant, and the varied pre-illumination experiments were performed on another plant. (B) Initial ΔΨ decay kinetics of the WT are shown (infiltrated in the dark with water or 50 mM TCEP), as well as the *gamera* mutant and the *ATPC1*-complemented *dpa1* mutant, termed *dpa1c.* The slowdown at ~100 ms is shown with an arrowhead. Dashed lines indicate the ΔΨ decay value 1 min after the pulse.

Fig 3A also shows the lifetime of 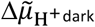 and ATP after illuminations of different durations (between 3-s and 12-min) in non-infiltrated leaves of the WT. Interestingly, if the duration of the illumination reached 7-s or more, the 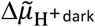 attained at the end of the illumination period became independent on the illumination (between ~-1.3 to −1.5 charge separations/PSI). However, a lag was observed before the 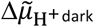 dark started to decay; whose duration increased with the time of illumination. Then, ATP levels decreased continuously after illumination and reached in ~ 20 min a stable dark level corresponding to the dark-adapted value (~-3.5 charge separations/PSI). In this view, the slow decrease of 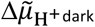 reflected the consumption of the ATP produced during the illumination or of the accumulated high energy bonds in different plastid metabolites, to which the ATP pool is buffered (see Discussion/Conclusion). The water-infiltration does not seem to have an impact on the behavior of the leaf, at least for illuminations not longer than 30 s. This is illustrated by the fact that the curve described before (30-s illumination of a water-infiltrated leaf) stands in between the 15-s and 40-s illumination curves for non-infiltrated leaves (Fig 3A). However, the behavior of the water-infiltrated leaves differ from the non-infiltrated ones for illuminations longer than 30 s, which might be due to the lower CO_2_ availability in water-infiltrated leaves, and therefore a lower activation of the Calvin-Benson cycle.

The final goal of this manuscript will be to establish the 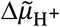 dependency of the CF_1_F_o_ activity in the reduced and oxidized forms. Accordingly, we compared leaves infiltrated with water or with TECP, which promotes γ-disulfide cleavage. We also looked at the behavior of the *gamera* mutant, in which the CF_1_F_o_ is in its reduced form (γ-dithiol) in the dark [33]. The 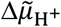 threshold 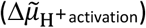 at ~100 ms in darkness, which is characteristic for water-infiltrated samples with the γ-disulfide (arrowhead in Fig 3B, see also Fig 1B), disappeared in presence of the TCEP due to accelerated 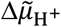 decay kinetics at the end of phase 2. Like in TCEP-infiltrated WT, the 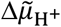 decay kinetics in the *gamera* mutant was accelerated and was even faster. In both cases, the 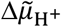 decay kinetics was completed in ~1 s. The faster decay kinetics in the *gamera* mutant, obtained in the *dpa1* genetic background [33], could be unambiguously attributed to the presence of the dithiol since the behavior of the *ATPC1*-complemented *dpa1* mutant, *dpa1c,* was similar to the WT (Fig 3B; ATPC1 is the thioredoxin-sensitive γ-subunit isoform and ATPC2 is the isoform present in *gamera).* The difference of ECS decay between *gamera* and TCEP-infiltrated WT leaves suggests that TCEP did only partially reduce the disulfide bond in the dark. Limited efficiency of TCEP in cleaving protein disulfides has been described elsewhere [47]. Based on the pronounced slowdown of the 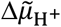 decay after phase 2, we conclude that CF_1_F_o_ was present mainly in its oxidized state in dark-adapted WT leaves, in contradiction with a previous report [16]. Addition of TCEP promoted the reduction of γ-subunit disulfide in a fraction of CF_1_F_o_ (distributed homogeneously along the thylakoid membrane), resulting in a mixture of active and inactive CF_1_F_o_ and a continuous decay kinetics of the 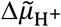.

In TCEP-treated leaves, the 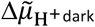 attained after the 30-s illumination was similar to the one in water-infiltrated leaves (~-1.5 charge separations/PSI, Fig 3A), but the lag was slightly shorter leading to a shorter lifetime of the 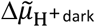 and ATP, respectively. We cannot rule out that TCEP infiltration also influenced the redox state of some Calvin-Benson cycle enzymes, altering the ATP sink capacity in the dark. Alternatively, TCEP could also produce mild uncoupling of the membranes. It is noteworthy, however, that at variance with the effect of commonly used dithiothreitol (DTT), the kinetics of the fluorescence increase in a dark-adapted leaf was not altered upon TCEP infiltration, eliminating a potential side effect on the PQ pool redox state (not shown). In a similar experiment, we analyzed the effect of a 14-min pre-illumination on a non-infiltrated *gamera* mutant leaf (closed circle in Fig 3A). At variance with the results obtained in the WT, we observed that illumination of the *gamera* mutant induced a decrease of the 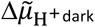 levels that returned to the dark-adapted level very slowly in ~ 45 min (see red horizontal line). We conclude that the mutation induced a perturbation of CF_1_F_o_ that prevented long term ATP accumulation during illumination. A lower *gamera* proton conductivity in steady-state light was reported [33] which might reflect this CF_1_F_o_ perturbation. The latter could be related to the large slowdown of Calvin-Benson cycle activation (about a factor 4) and the larger transient NPQ formation observed on the mutant with respect to the WT (see Fig S5).

### 4. 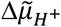 dependence of CF_1_F_o_ rates in dependence of the γ-subunit redox state

The final aim of this work is to determine how the reduction/oxidation of the γ-disulfide modifies the 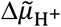 regulation of CF_1_F_o_ activity. To modulate the redox state of the γ-disulfide, we compared dark-adapted water-infiltrated leaves (oxidized form) with TCEP-infiltrated leaves (partially reduced form) and the *gamera* mutant (fully reduced form). As discussed in the introduction, studying the 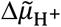 regulation of CF_1_F_o_ activity requires the *in vivo* measurement of both the 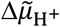 and the CF_1_F_o_ activity. In this work, the pulse-induced ECS method to measure 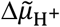 has been established and validated in both redox states. For CF_1_F_o_ activity determination, the ECS decay rate after a single turnover laser flash is routinely used [e.g., Ref. 40]. However, there is a bias due to the “*b* phase”, a ~10-ms ECS rise due to electrogenic contribution of the cytochrome *b_6_f* activity which superimposes to the CF_1_F_o_ activity-related ECS decay. Moreover, we observed that another source of variation occurred when using flashes, which has been documented already [10, 11] but remains poorly understood. We tried to circumvent these sources of error and mitigate their contributions to the ECS decay by generating the 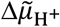-activated state of CF_1_F_o_. The latter was routinely obtained by short saturating pulses which had a duration of at least 8-ms (and lasted up to 17-ms depending on the 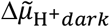; see Fig S6). We also measured the rate of 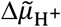-activated CF_1_F_o_ as the slope of the ECS decay when ECS signals reached the level expected after a saturating laser flash (i.e., 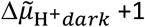 charge separation/PSI, see Fig S6). At that time, the cytochrome *b*_6_*f* activity was mostly over and/or contributed marginally to the apparent kinetics: the *b*_6_*f*-catalyzed electron transfer between P700^+^ and PQH2 lasted longer than after a flash but this process was negligible at the time of the measurement compared to the coincident ΔΨ consumption by 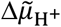-activated CF_1_F_o_.

Fig 4A sums up all the CF_1_F_o_ rates for various 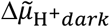 values that were measured with this method. All data presented in this manuscript so far have been included, i.e. in WT leaves (dark adapted or during relaxation following a 30-s illumination), in TCEP-treated WT leaves in the same conditions, as well as in *gamera* (dark-adapted or following a 14-min illumination). The dark-adapted situation in the presence of AA was also added. As discussed before, the *gamera* mutant represents the behavior of the reduced form of the CF_1_F_o_ whereas the water-infiltrated WT represents its oxidized form and the TCEP-infiltrated WT is an intermediate situation where part of CF_1_F_o_ is in the oxidized form and part of CF_1_F_o_ is in the reduced form. Below 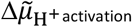 (−2.2 charge separations/PSI, see arrow in Fig 1B, Fig 3B), the CF_1_F_o_ was always faster in the reduced form (TCEP-infiltrated or *gamera)* than in the oxidized form (water-infiltrated). In both reduced and oxidized forms, the CF_1_F_o_ rate vs. 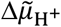 curves resembled the ones obtained *in vitro* by Junesch and Gräber [36]. The rate of CF_1_F_o_ increased exponentially with 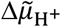 in the reduced form, whereas a lag was present in the oxidized form until a specific 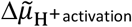 was reached.

**Fig 4:**
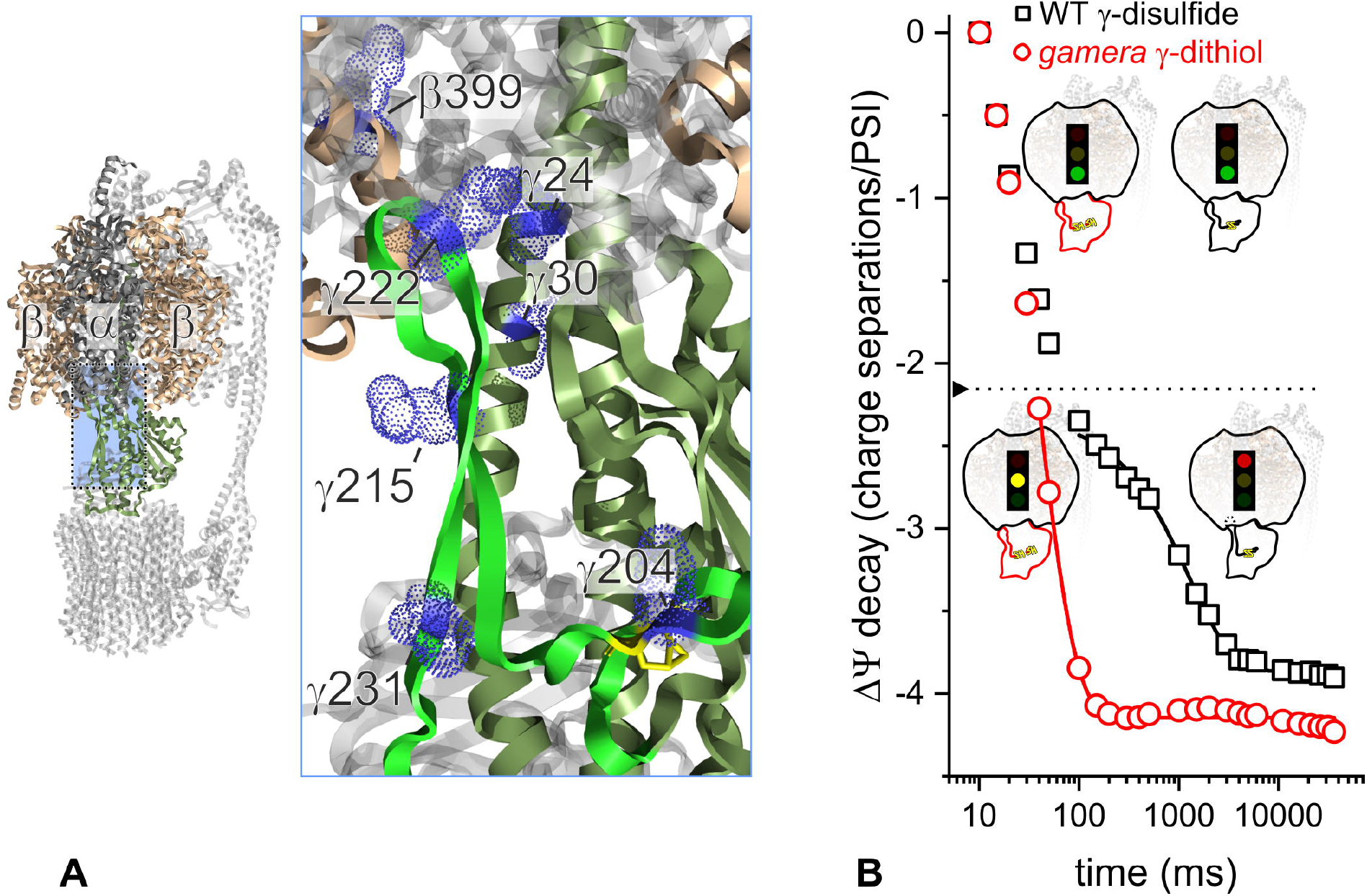
The regulation levels of 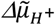 and thiol modulation on ATP synthesis are shown. (A) The 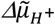-dependence of the CF_1_F_o_ rate in the oxidized (dark) and reduced (red) forms are shown, as well as in the intermediate situation of TCEP-infiltrated WT leaves (blue). Rates were obtained according to Fig S6. Closed symbols: untreated leaves, open symbols: leaves were additionally infiltrated with 40 μM antimycin-A (+AA). Fitted curves were obtained with Equation 1. The water-infiltrated WT showed a 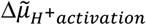 of ~-2.2 charge separations/PSI, and half-maximal *R* at ~-1.5 charge separations/PSI (dotted black line). The γ-dithiol-containing *gamera* showed a half-maximal *R* at ~-3.8 charge separations/PSI (dotted red line) and a a 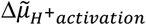 of ~-4.7 charge separations/PSI. (B) The decay rate *R* (see legend, in s^-1^) of the CF_1_F_o_ was calculated in the range from 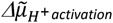 (arrowhead) to 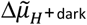 levels. The relaxation of ECS was fitted by a mono-exponential decay function.

Both 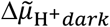-dependent rates of (chemically unmodified) CF_1_F_o_ in WT and *gamera* have been fitted with a sigmoid function, like in [36]. In fact, the plotted CF_1_F_o_ rates of both genotypes were processed in one calculation by using a sum of two sigmoid functions. In Equation 1, we forced, via the constant *c*, the first summand to be zero in the *gamera* plot and the second summand to be zero in the WT plot:

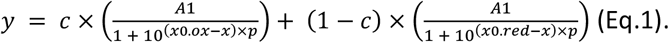

*A1* defines shared maximal rates. With the respective standard errors, *A1* was 35.3 ± 1.1 s’^1^. The slope parameter of the function is expressed as *p* (1.4 ± 0.1). The 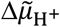 where the half-maximal CF_1_F_o_ rates were obtained for WT and *gamera* are defined as *x0.ox* (−1.5 charge separations/PSI) and *x0.red* (−3.8 charge separations/PSI), respectively. The sigmoid TCEP curve in Fig 4A was obtained with *c* = 0.56 ± 0.04, suggesting about 56% of CF_1_F_o_ in the oxidized form (γ-disulfide conformation) upon infiltration with TCEP. This value was close to the fraction of reduced CF_1_F_o_ in the presence of TCEP based on the kinetics of ECS decay in the *gamera* and WT presented in Fig 4B (already presented in Fig 3B). In Fig 4B, the relaxation of ECS from the 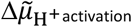 (arrowhead) to 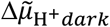 (~-4 charge separation/PSI for the 3 curves) was fitted with a mono-exponential decay. Rate constants of 0.8 s^-1^ in water-infiltrated WT were obtained in the presence of ~100% γ-disulfide, and 30.6 s^-1^ were measured with 100% γ-dithiol in *gamera.* Accordingly, Fig 4B suggests that TCEP infiltration produced ~58% γ-disulfide cleavage in WT CF_1_F_o_, since the rate becomes 13.4 s^-1^.

In line with previous reports on high ATP synthesis rates under disulfide-promoting conditions [22, 36], we did not observe an effect of the redox regulation on the maximal CF_1_F_o_ rate at high 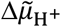 in WT, even though we did not manage to reach high enough 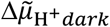 to fully describe the plateau of CF_1_F_o_ rate. This was due to experimental constraints: in the oxidized form, it was not possible to reach higher values of 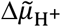 because a sufficient dark period was needed after the 30 s illumination to reoxidize PSI secondary donors and PSII acceptors, as discussed before. In the reduced CF_1_F_o_ of the *gamera* mutant, illumination decreased the 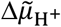 (Fig 3A). However, the observation that TCEP-(~50% γ-dithiol) and water-infiltrated WT leaves reached similar rates at 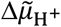 ~-0.5 charge separation/PSI (see also Fig S6, panel A) suggests that the rate of the oxidized and reduced form reached similar values in this range of 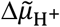.

## Conclusion/ Discussion

Taken together, the *in vivo* ECS protocol that was developed previously [16] allowed to estimate the 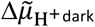 by avoiding formation of a ΔpH during probing of ΔΨ. The 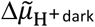 dark is defined by the [ATP]/([ADP][Pi]) ratio and was maintained at high levels after a photophosphorylation period. It also required metabolic coupling with mitochondria which were most efficiently inhibited by uncouplers. By comparing the *gamera* mutant with WT, the 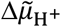-dependent CF_1_F_o_ activity profile demonstrated higher efficiency of ATP synthesis in the reduced form at lower membrane energization levels. In this work, we could confirm the redox regulation of the CF_1_F_o_ previously described by Junesch and Gräber *in vitro* [36] in which a chemically reduced disulfide bond in the γ-subunit produced half-maximal rates at a lower 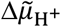 of 0.7 ΔpH units. In our work, we obtained a very good agreement with this value since we measured that the 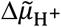 required for half saturation of the CF_1_F_o_ was 2.3 charge separations/PSI higher in the oxidized form (WT) compared to the reduced form *(gamera* mutant). Accepting a value of 20 mV per PSI charge separation [38, 48, 49], this determined value (~45 mV) was very close to the 0.7 pH units.

Although the pH shift between the two sigmoids in the *in vitro* work of Junesch and Gräber, and the one we obtained *in vivo* were in good agreement, our results do not allow to strictly compare the absolute values of the electrochemical gradient needed to reach half-saturation. This comes from the fact that the 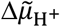 values in our work are expressed in relative values. For a proper *in vitro* and *in vivo* comparison, our results should be expressed in absolute values, i.e. compared to the situation where 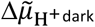 dark is null.

### How to reach the situation where 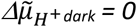 ?

Coming back to our opaque bottle analogy, the water volume contained in an opaque bottle can be easily calculated as the difference between the empty volume of the bottle (measured as the volume of water that needs to be added before it spills out) and the maximal countenance capacity (same measurement with previously emptied bottle). Several studies [16, 46] have shown that the electrochemical proton gradient in the dark 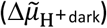 can be collapsed by infiltrating the samples with appropriate inhibitors. This is the equivalent of the “emptied bottle” situation, and allows expressing the values of 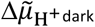 as absolute values, like in the work of Junesch and Gräber, instead of relative values (by comparison to ECS_10ms_). In brief, the ECS-based protocol to measure the absolute value of 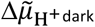 is the following: 1-energizing the membrane with a very strong light pulse and probing the ECS increase until the 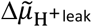 is reached (i.e., measuring the “empty volume”), 2-collapsing the 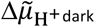 with appropriate inhibitors, and 3-repeating the experiment starting from such a situation where 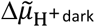 is null (i.e., measuring the “maximal countenance capacity”). The difference between the amplitudes of the two pulse-induced ECS increases corresponds to the absolute 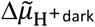 in the non-treated sample. To do this, one needs to find a set of inhibitors that totally collapse the 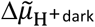. The latter is built at the expense of the ATP present in the plastid in dark-adapted leaves, which originates from mitochondrial ATP production as has been shown previously for algae [50], diatoms [46], and higher plants [16]. In mature plant tissues, the cytosolic ATP import is relatively slow due to low abundance of nucleotide transporters in the chloroplast membrane [45]. Previous reports [16, 46] and the first section of this manuscript used antimycin-A (AA) treatment, which inhibits the bc1 complex in the mitochondrial respiration. This was proposed to suppress mitochondrial ATP production and therefore to collapse 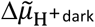 [16]. However, it is known that in the absence of bc1 activity, a mitochondrial electron flow can still occur thanks to the activity of the alternative oxidase (AOX). Despite its non-electrogenicity, the AOX allows the complex I to function and to generate an electrochemical proton gradient and mitochondrial ATP. In agreement with this, the use of AA alone does not fully collapse the electrochemical gradient in diatoms [46] and only the inhibition of both the bc1 complex (with AA) and the AOX (with salicylhydroxamic acid) fully suppressed the electrochemical gradient. To determine the best protocol to collapse the 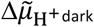 in plants, we compared different pharmacological treatments (Fig S4). Fig S4 displays the ECS decay kinetics after a pulse in dark-adapted leaves treated with various inhibitors that affect the 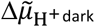. We used two treatments which should fully collapse the 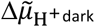: Nonactin is a specific K^+^ transporter which in combination with Nig should remove both the ΔΨ and ΔpH across the thylakoid in the dark, as should FCCP, which is an uncoupler. The dark-adapted values of the 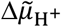 were very similar between the two treatments as well as with the treatment with Nig alone, indicating that Nig was sufficient to suppress the mitochondrial ATP production and, in turn, the 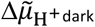. However, it is important to note that the pulse-induced 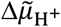 had a very long lifetime (~45 – 60 s) in Nig-infiltrated leaves. This reflected, in the absence of mitochondrial ATP supply, the interplay of Nig-dependent ΔpH/ΔΨ exchange and the consumption of the ATP generated during the pulse. We then compared AA treatment to the Nig treatment and obtained variable results in different plants. In some cases (Fig S4B), the dark-adapted value of the 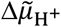 in AA was very similar to the one in Nig, indicating that AA inhibited most of the mitochondrial ATP production whereas in other cases (Fig S4C) Nig and AA treatments gave very different results, suggesting that the activity of the AOX was still significant. To conclude, when one wants to express 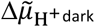 in absolute values (i.e. to measure the “total volume” of the opaque bottle), the reference for 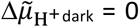 should be obtained with Nig, which suppresses mitochondrial ATP production, or with thylakoid uncouplers collapsing the 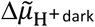 across the thylakoid (e.g. FCCP or the combination of Nig and nonactin). However, it is important to note that since these compounds embedded to the lipid bilayer, fast and slow ECS decay kinetics will be altered and cannot be attributed to CF_1_F_o_ performance alone.

With this in mind, it is possible to discuss our results in the light of the 2.7 and 3.4 pH units for half saturation of the reduced and oxidized ATP synthesis rate *in vitro* [36], which would correspond here to 162 and 204 mV. With a calibration of 20 mV per PSI charge separation, this would give ~8.1 and ~10.2 charge separations/PSI. For the reduced and oxidized ATP synthesis rate *in vivo,* the half-saturation was reached at a 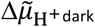 of −3.8 and −1.5 charge separation/PSI. Here, the lowest value of 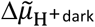 was obtained for AA-infiltrated WT leaves in the presence of TCEP, which was ~-7.5 charge separations/PSI (note that *x*-axis in Fig 4A is 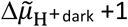). As outlined above, the 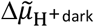 in TCEP/AA samples was significantly higher than 0. In fact, the remaining 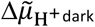 in TCEP/AA should be at least 4 charge separations/PSI to explain the discrepancy (4.2 = 10.2 – 7.5 + 1.5 charge separations/PSI; 84 mV or 1.4 pH units for oxidized CF_1_F_o_). Such a value being high according to the results of Fig S4, we acknowledge here a slight discrepancy between our results and the ones obtained *in vitro* [36].

### What does the lifetime of ATP depend on?

In this work, we could measure the lifetime of ATP after an illumination and showed in section 3 that the ATP accumulated after an illumination saturates after 7 s of illumination, and that longer illumination durations lead to a lag before the ATP pool relaxes. We favor the hypothesis that the increased ATP lifetime (closed squares in Fig 3A), as a function of the pre-illumination period, was associated with the accumulation of high energy bonds in different plastid metabolites. This hypothesis is supported in a nucleotide transporter study which revealed that the ATP exchange rate between chloroplast and cytosol was rather slow, and fluorescent ATP probes suggested that excess of free ATP during illumination diminished rapidly in the dark [45]. Thus, lowering the 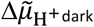 in Fig 3A would be delayed after prolonged illumination due to relatively slow metabolite conversions that consume ATP in the dark. For instance, high levels of ribulose-1,5-bisphosphate and 1,3-bisphosphoglycerate in the Calvin-Benson cycle could be part of a feedback inhibition that prevents their own ATP-consuming synthesis. Furthermore, the deactivation of the Calvin-Benson cycle in the dark [16] was in the same time range as the decay of 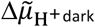. Thus, ATP is stored as 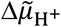 and, instead of leak processes, slow metabolic consumption occurs within the chloroplast. When infiltrating leaves with TCEP (midpoint redox potential of −290 mV [51]), various thiol enzymes are expected to be partially reduced due to the chemical which could lead to their activation. Usually catalyzed by reduced thioredoxin, TCEP could activate Calvin-Benson cycle enzymes such as fructose-1,6-bisphosphatase or sedoheptulose-1,7-bisphosphatase. This, in turn, could explain a slightly earlier onset of the consumption of ATP after 30-s illumination in TCEP-infiltrated samples (cf. open squares in Fig 3A), which we measured on the 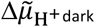 levels.

### Is the dark-adapted CF_1_F_o_ fully oxidized?

In the fit of the water-infiltrated experimental data (black curve, Fig 4A), we considered that the dark-adapted CF_1_F_o_ was fully oxidized. Nevertheless, about 5% of CF_1_F_o_ were reduced in a previous study of dark-adapted spinach chloroplasts [2], suggesting that γ-dithiols in water-infiltrated samples were likely not fully oxidized. Also, the rate we measured in the low 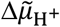 range (Fig 4B) was not null although it was previously proposed that the CF_1_F_o_ was fully inactive below the 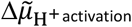 [10, 11]. If so, the rate constants measured in Fig 4B (0.8 s^-1^ in control vs. 30.6 s^-1^ in *gamera)* would mean that in our case, 2-3 % of the CF_1_F_o_ is in the reduced form in the dark-adapted state. At 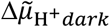 levels below the 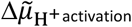, the contribution of ion channel activity to the ECS decay was likely more significant in WT leaves.

### Uncoupler effect in TCEP-infiltrated leaves?

In Fig 4A, the fit of the TCEP-infiltrated experimental data by the sum of two sigmoids failed for the AA-treated samples: significantly elevated rates were observed when the 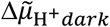 was low in the presence of TCEP + AA (open blue diamonds). We cannot rule out that TCEP infiltration had a weak uncoupling effect or modified anion channel activity, thus producing slightly faster ΔΨ decay rates. This effect would slightly overestimate the TCEP-dependent rates in Fig 4A. In line with this, the slightly earlier decrease of 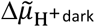 in TCEP-infiltrated samples upon illumination might be due to a slow leak process (Fig 3A). The leak might be due to activation of anion channels or deactivation of H^+^/cation antiporters. It has also been reported that DTT treatment of chloroform-extracted CF1 resulted in the reversible dissociation of the ε-subunit [52, 53] which, if this is the case for CF_1_F_o_ *in vivo,* would produce uncoupling [54]. It remains to be tested whether the elevated ECS decay rates in the presence of TCEP, most pronounced at low 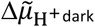 levels (open blue diamonds in Fig 4A), were due to ion channels other than CF_1_F_o_.

### Combining function, structure and physiology

In the light of the recently obtained structure of the CF_1_F_o_, we would like to try linking our results to structural considerations. The L-shaped double hairpin (Fig 5A, light green in magnified area) in the disulfide-containing spinach CF_1_F_o_ structure interacts with the β-subunit DELSEED motif and has therefore been suggested to prevent ATP hydrolysis by forming a chock during rotation [3]. Steric interference of rotation in the opposite direction (ATP synthesis) can thus be proposed for this structural element as well. The γ-hairpin and the β-DELSEED loop harbor various residues that are not surface-exposed (e.g. γK222 and β399 in Fig 5A) but have been chemically modified in illuminated thylakoids only [27, 28, reviewed in 29]. Thus, the structural impact of the 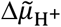 on CF_1_F_o_ is occurring (among other sites) in the rotational chock element. Solvent exposure of the latter should reduce the bulk of rotating γ-residues in 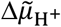-activated CF_1_F_o_ which modify the sum of van der Waals contacts with the β-DELSEED loop. This β-subunit loop moves perpendicular to the γ-subunit axis and the stroke movements (exerted by γ and β during ATP synthesis and hydrolysis, respectively) eventually change the binding site conformation and thus the nucleotide affinity [55]. The sequential binding site cooperativity is most efficient in 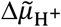-activated CF_1_F_o_, yielding maximal rates of the enzyme in both redox states (Fig 5B above dashed line). When the 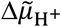 approaches lower levels, the choc structure becomes buried again and rotational catalysis occurs at lower efficiency (Fig 5B WT below dashed line). The L-shaped double hairpin position is stabilized by the γ-disulfide (Fig 5A, yellow in magnified area) which might explain why the reduced enzyme remains in a highly active state at lower 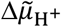 (Fig 5B *gamera* below dashed line). From a physiological viewpoint, enabling higher CF_1_F_o_ rates for a given 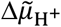 level upon γ-disulfide cleavage has the advantage to maintain a high H^+^ conductivity via CF_1_F_o_ even in sub-saturating light conditions. Thus, ATP production is maximized and electron transfer limitation (triggered by an acidic lumen for instance) can be avoided. Probably therefore, disulfide cleavage *in vivo* is very efficiently realized in dim light [19, 20] and at higher light intensities [21–23, 28, 56].

**Fig 5:**
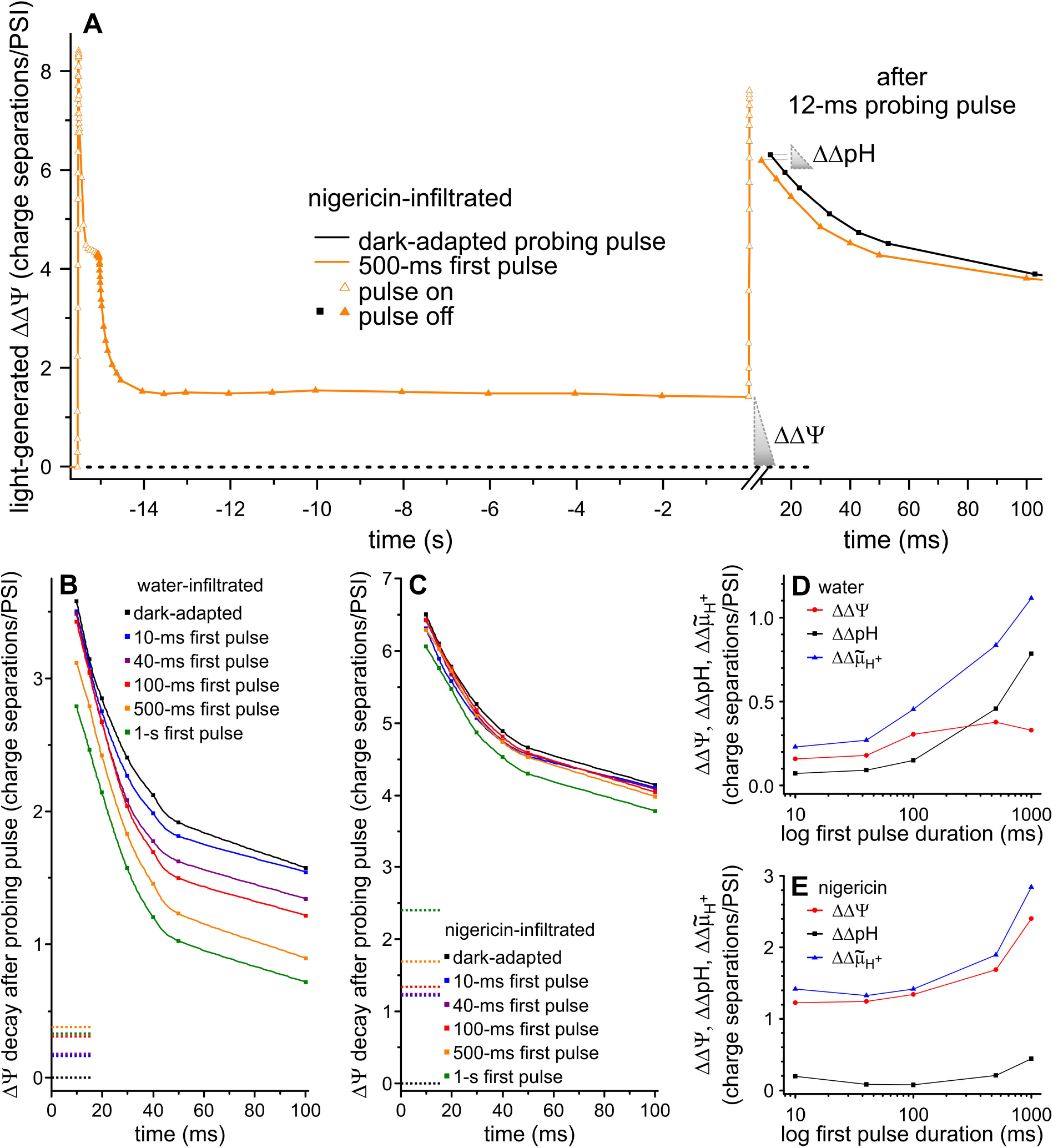
A selection of 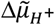-dependent structural rearrangements in spinach CF_1_F_o_ and a tentative model for different ATP synthesis rates as a function of 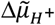 are shown. (A) The structure of spinach CF_1_F_o_ (PDB ID 6fkh) magnifies the redox loop (light green, disulfide in yellow sticks) on the right. The frontal α-subunit is magnified half-transparent for the sake of visibility of the γ-subunit (dark green). The ATPase choc structure is formed by the γ-hairpin loop around position 222 which is stabilized by the disulfide [3]. A variety of structural rearrangements are expected depending on the 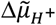, including the γ-hairpin. Critical residues (blue spheres) become available for chemical labeling or trypsin digestion exclusively in illuminated thylakoids [reviewed in 29]. (B) The exposed position of the choc structure in 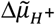-activated CF_1_F_o_ allows efficient rotational catalysis in both γ-redox states (top). Below the WT activation threshold (dashed line), the disulfide-stabilized choc structure prevents efficient ATP synthesis. The *gamera* sustains an active conformation at this 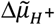 since the choc structure is less efficiently fixed inside the α_3_β_3_ cavity, owing to the absence of the γ-disulfide.

## Supporting information

Supplemental Material combined

## Acknowledgements

We would like to thank Dr. Jörg Meurer (Ludwig Maximilian University of Munich) for sharing the dpa1c and *gamera* seeds with us. B.B. and F.B. acknowledge funding from the ERC Starting Grant PhotoPHYTOMICS (ERC-2016-STG grant # 715579). F.B. also acknowledge the CNRS and the “Initiative d’ Excellence” Program from the French state grant “DYNAMO”, ANR-11-LABX-0011-01.

## Notes

### Competing Interest Statement

The authors have declared no competing interest.

