## Supplemental Material combined for "ECS-based investigation of chloroplast ATP synthase regulation"

Supporting Information – Supplementary Figures S1-S6

Supplemental reference list

Supplementary Figure 1

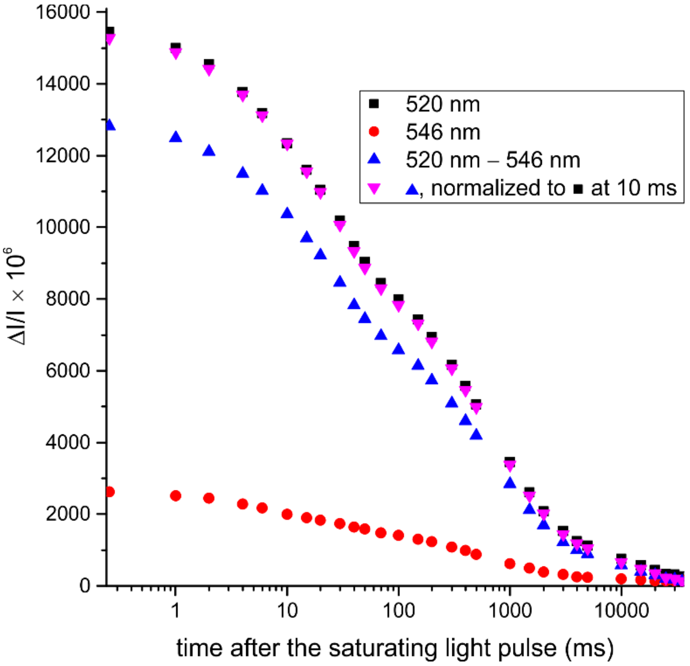


Figure S1: Decay of electrochromic shift signals measured at 520 nm (dark squares) and 546 nm (red circles), respectively, after a 12-ms saturating light pulse on a logarithmic time scale. The differential signals is shown in blue triangles and, given that the $\Delta$I/I level at 10 ms darkness served as a reference for our study, is normalized to the 520 nm at 10-ms (magenta downward triangle). The very minor deviations in the kinetics (magenta vs dark) supported the validity of single-wavelength measurements when using short pulses in studies of the $\Delta\tilde{\mu}_{\mathrm{H^{+}}\,\mathrm{dark}}$ dynamics.

Supplementary Figure 2

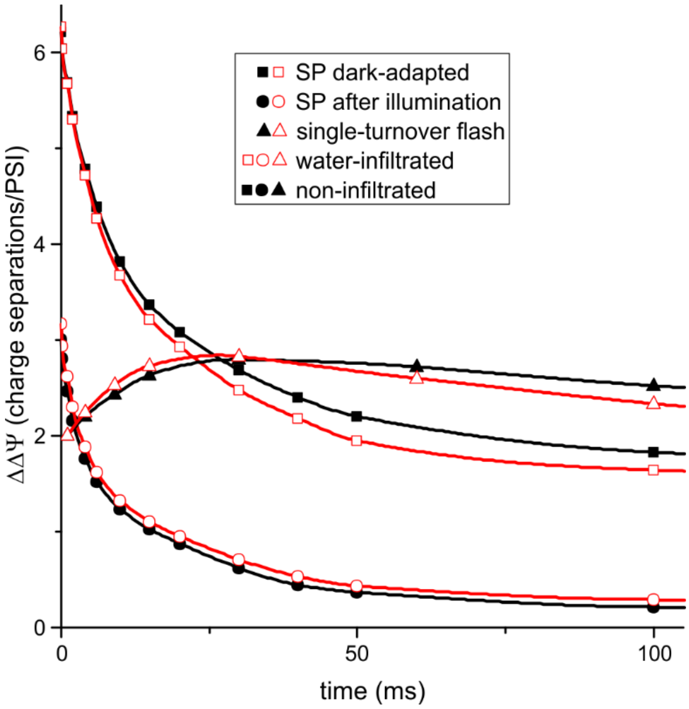


Figure S2: Conservation of physiological properties upon infiltration of leaf samples. ΔI/I signals dropped by ~30% upon infiltration due to longer light trajectory in non-infiltrated samples. However, once the signals were normalized to the ECS increase following a saturating laser flash (see Methods), the behavior with (open red) and without (close dark) water infiltration were very similar, following a single turnover flash (triangles) or a saturating pulse in fully dark-adapted states (squares). This was also true after applying a saturating pulse to samples that were pre-illuminated for 30-s and kept in the dark for 3-min (circles).

Supplementary Figure 3

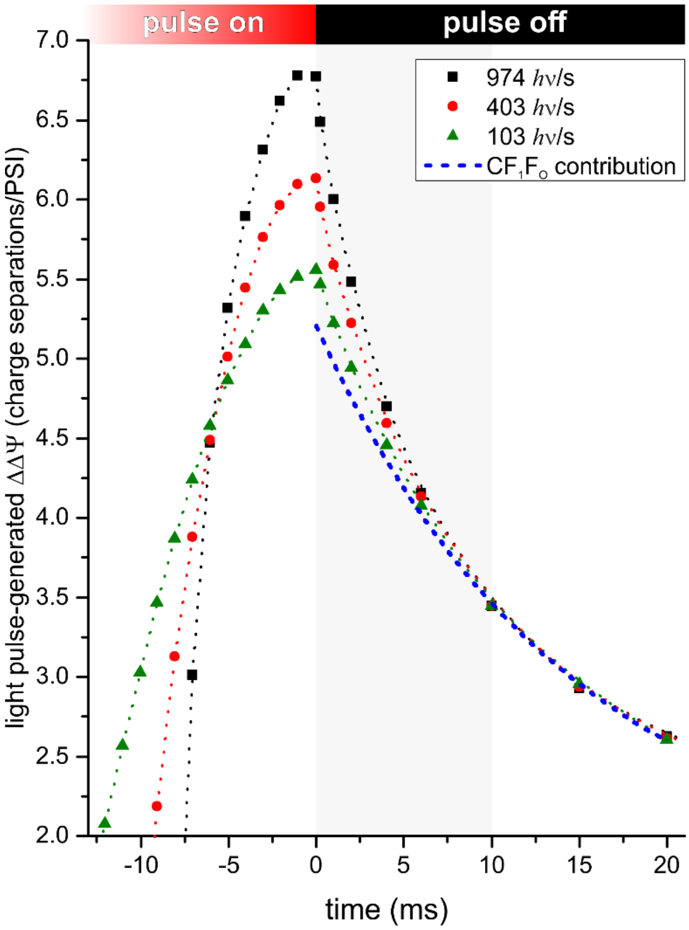


Figure S3: Analysis of the initial decay kinetics upon generating a maximum membrane potential by light pulses of various intensity. The ECS decay rates in the first 10 ms of darkness (gray area, termed leak phase) depended on the intensity of the light pulse. We assume that the plateau value was determined by $H^+$ leaks that were triggered above a critical $\Delta\tilde{\mu}_{H^+}$ potential ($\Delta\tilde{\mu}_{H^+ \text{leak}}$). Above this potential, the rate of ATP synthesis at the level of the $F_1$ complex might become a rate limiting process, inducing temporary decoupling of $H^+$ translocation from ATP synthesis. As shown previously for spinach [cf. Fig. 1 in supplemental reference 1], the decay kinetics after 10 ms were identical and reflected $H^+$ translocation via $CF_1F_o$.

Supplementary Figure 4

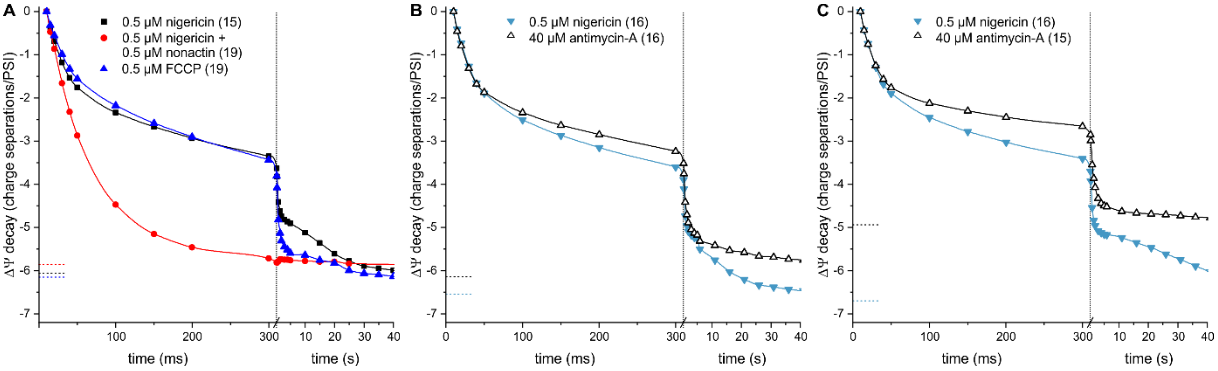


Figure S4: The effect of mitochondrial inhibitors and uncouplers on the $\Delta\tilde{\mu}_{H^+ \text{dark}}$. (A) Comparison of the pulse-induced ECS kinetics in three treatments, nigericin (black squares), nigericin + nonactin (red circles) and FCCP (blue triangles) demonstrate similar $\Delta\tilde{\mu}_{H^+ \text{dark}}$. The three curves were obtained on leaves from the same plant and digits in parenthesis indicate the duration of the pulse in ms. When the $ECS_{10ms}$ is set to 0, the $\Delta\tilde{\mu}_{H^+ \text{dark}}$ is visualized as dashed lines, which is the ECS value obtained after ~1 min darkness. (B) Representative example of comparison between antimycin-A and nigericin treatment on leaves from the same plant. The similar values of $\Delta\tilde{\mu}_{H^+ \text{dark}}$ indicate a low AOX activity. (C) Representative example of comparison between antimycin-A and nigericin treatment on leaves from another plant. The very high $\Delta\tilde{\mu}_{H^+ \text{dark}}$ in AA-infiltrated leaf indicates a high AOX activity.

Supplementary Figure 5

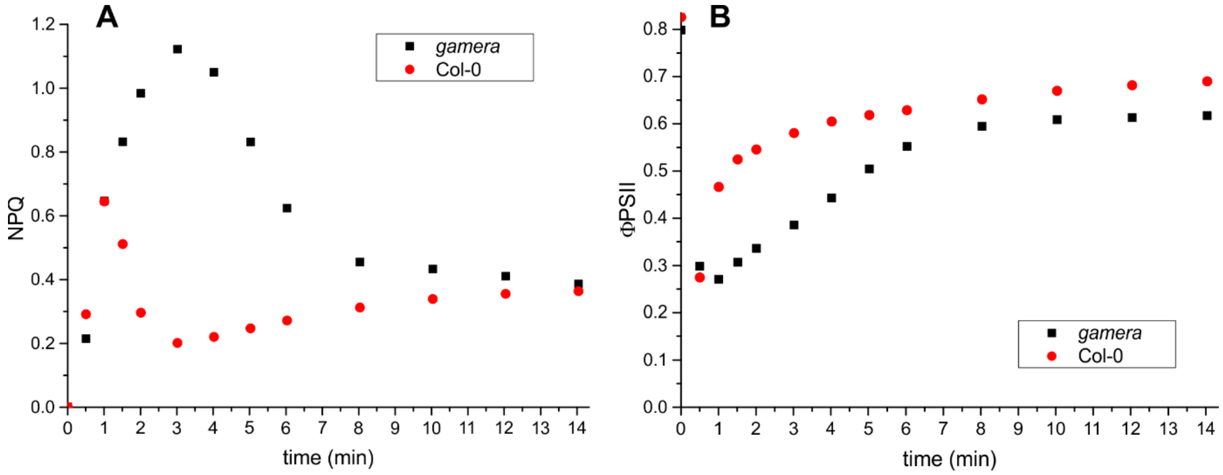


Figure S5: Typical kinetics of (A) NPQ and (B) ΦPSII following a dark-to-light transition in leaves of Arabidopsis wild type (red) and *gamera* mutant (black). During illumination of the mutant, a prolonged build-up of NPQ and a delayed activation of photosynthesis was observed.

Supplementary Figure 6

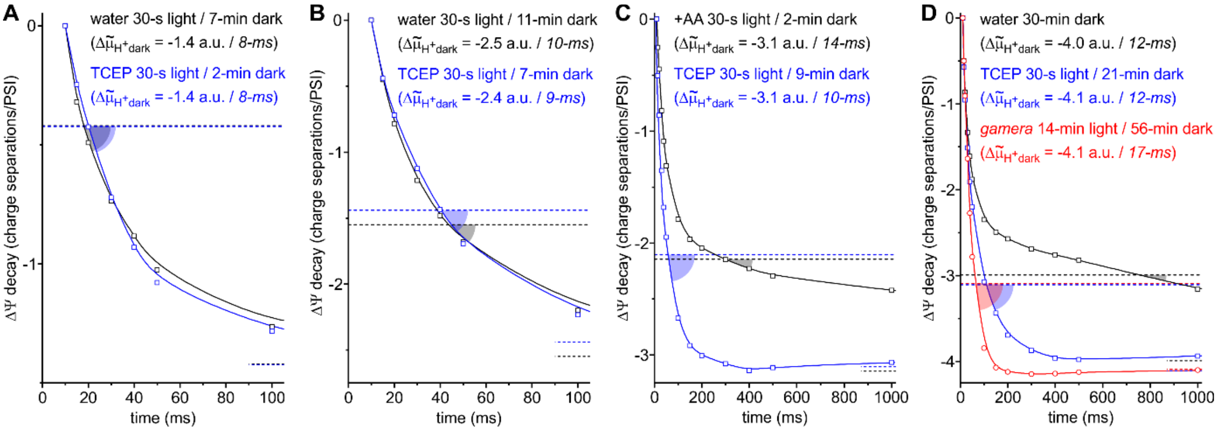


Figure S6: Calculation of the $CF_1F_o$ rate from the ECS decay kinetics is shown in water-, TCEP-infiltrated WT and *gamera* (only in panel D since *gamera* could not accumulate ATP in the light and therefore could not sustain a high $\Delta\tilde{\mu}_{H^+dark}$). The data is used in Fig 4A of the main text. The rationale was to calculate the rate of the $CF_1F_o$ when the electrochemical proton gradient reached $\Delta\tilde{\mu}_{H^+dark}$ +1, to relate to the classical protocol based on the ECS decay following a single turnover flash. Each panel compares samples with similar $\Delta\tilde{\mu}_{H^+dark}$ (-1.4, -2.5, -3.1 and -4 charge separations/PSI) from Fig 1B/D and Fig 3A in the main text. The dashed lines on the bottom right indicate the $\Delta\tilde{\mu}_{H^+dark}$ and the full-width dashed lines represent $\Delta\tilde{\mu}_{H^+dark}$ +1 charge separation/PSI. The semi-transparent angles visualize the ECS decay rate at $\Delta\tilde{\mu}_{H^+dark}$ +1 charge separation/PSI, expressed as *R* in Fig 4A. The $\Delta\tilde{\mu}_{H^+dark}$ was estimated by a ms-pulse and its duration is given in italics. In panels A and B, the $\Delta\tilde{\mu}_{H^+dark}$ remained above the $\Delta\tilde{\mu}_{H^+activation}$ and only a fast phase of ECS decay was seen in both γ-redox states. In panels C and D, the $\Delta\tilde{\mu}_{H^+dark}$ was below the $\Delta\tilde{\mu}_{H^+activation}$, and the kinetics of ECS decay in the water-infiltrated leaf became biphasic. As discussed in Sections 1 and 3, the break observed in the $\Delta\tilde{\mu}_{H^+}$ decay kinetics at the end of phase 2 revealed a $CF_1F_o$ transition from an active toward an inactive form, which was only observed under γ-disulfide promoting conditions (oxidized, water-infiltrated WT).
